# Integrative systems neuroimmunology reveals leukocyte-expressing PAX6 as a critical predictor of major depressive disorder

**DOI:** 10.1101/2024.09.25.614771

**Authors:** Haroldo Dutra Dias, Anny Silva Adri, Adriel Leal Nóbile, Marilia Garcia de Oliveira, Elizabeth N. Chung, Ian Antunes Ferreira Bahia, Dennyson Leandro M Fonseca, Lena F. Schimke, Igor Salerno Filgueiras, Pedro Marçal, Fernando Yuri Nery do Vale, Rodrigo J S Dalmolin, Gustavo Cabral-Miranda, Helder Nakaya, Renato Bortoloti, Clement Hamani, Michael A. Wheeler, Rafael Machado Rezende, Otavio Cabral-Marques

## Abstract

Major depressive disorder (MDD) is a complex psychiatric condition with a significant global impact. This study applied a genomic-driven integrative systems neuroimmunology approach to analyze transcriptomic data from 3,114 individuals (1,877 MDD patients and 1,237 controls). The analysis revealed neuroimmunological transcriptomic alterations, indicating cross-talk between the immune and nervous systems in peripheral blood mononuclear cells (PBMCs) and specific brain regions. Among 31 shared genes, NEGR1, PPP6C, SORCS3, and PAX6 emerged as significant predictors of MDD in patients’ PBMCs. Notably, PAX6 was also identified as a differentially expressed gene (DEG) in the amygdala, while NEGR1, PPP6C, and SORCS3 showed no significant differential expression in other central nervous system (CNS) regions. Validation by immunophenotyping in a mouse model of chronic stress demonstrated increased PAX6 expression in PBMCs, a gene previously associated with MDD in GWAS studies. Collectively, our findings suggest the existence of shared transcriptomic modules across the brain and immune system, highlighting PAX6 as a potential therapeutic target in MDD.

## INTRODUCTION

Major depressive disorder (MDD) is one of the most pressing and pervasive mental health challenges worldwide. It is characterized by apathy^1^, loss of interest or pleasure in previously enjoyable activities, suicidal ideation, cognitive impairment, neuropsychiatric comorbidities^1^, and painful physical symptoms (PPS)^2^. MDD results in years lived with disability and has become one of the leading causes of premature mortality^3^. Approximately 300 million people globally are affected by MDD, and its incidence is increasing annually, resulting in significant socioeconomic burdens^4^. Despite its prevalence, the pathophysiological mechanisms underlying MDD remain incompletely understood, posing tremendous hurdles to the development of effective treatments^5^.

In recent decades, advancements in genomics and transcriptomics have opened new avenues for understanding the neuroimmunobiological underpinnings of this complex disorder. Genomic studies, particularly genome-wide association studies (GWAS)^6^, have identified numerous genetic variants associated with an increased risk of MDD. These studies have shed light on the polygenic nature of MDD, implicating a wide array of genes involved in neurological pathways. However, despite these findings, the relationship between these genes, their expression profiles, and functional validation through immunophenotyping remains inconsistent. This gap highlights the need to identify reliable biomarkers and therapeutic targets for MDD.

Transcriptomic analyses have shed light on gene expression changes in both the brains^7^ and immune cells^8^ of individuals with MDD. While the brain transcriptome is critical for understanding MDD, its study is limited by the availability of postmortem data, which restricts insights into its dynamic nature during the course of the disorder. Therefore, integrating transcriptomic data from both the brain and immune cells of MDD patients, alongside genome-wide bioinformatic analyses, is essential^9^ for advancing our understanding of MDD neuroimmunology.

Neuroimmunology explores interactions between the nervous and immune systems, and is an emerging research area that has amassed evidence from both MDD patients and animal studies^10^. This reveals a bidirectional interaction between the nervous and immune systems. This interaction influences an individual’s stress response and significantly contributes to their susceptibility to mood disorders^10,11^. Emerging evidence indicates that neuroimmune processes play a pivotal role in MDD, where immune dysregulation and inflammation are recognized as central components of its pathogenesis. This has led to the development of the neuroimmune network model of depression^12^.

This study, using an integrative systems approach, characterizes an immunological network within the leukocytes of MDD patients, including the stress-induced expression of PAX6. For the first time, we demonstrate that PAX6, suggested in prior GWAS studies as potentially associated with MDD, is transcriptionally, at the protein level, and functionally linked to MDD. This is evidenced by the stress-induced expression of PAX6 in both human MDD patient samples and in the peripheral blood mononuclear cells (PBMCs) of mice subjected to chronic restraint stress (CRS). Our findings suggest that alterations in PAX6 expression in leukocytes, a transcription factor traditionally linked to neurodevelopment and brain function, reflect a broader connection between the brain and the peripheral immune system. By exploring these interactions, our study uncovers potential biomarkers and novel therapeutic pathways for MDD intervention.

## RESULTS

### Clusters of neuroimmunological biological processes in MDD transcriptomes

To identify the shared differentially expressed genes (DEG) and biological processes (BPs) from MDD patients across multiple transcriptome datasets from MDD patients (demographics, diagnosis, and treatment information available in **Supplementary Table 1**), we performed an integrative analysis. This approach aimed to uncover shared molecular mechanisms between PBMCs and the anterior cingulate cortex (ACC) by utilizing primary datasets publicly available at the Gene Expression Omnibus (GEO) repository. **Extended Data Fig. 1** provides an overview of primary data sets containing the total number of transcriptomes (3,114 individuals: 1,877 MDD patients and 1,237 controls) and the bioinformatics workflow.

We performed a differential expression analysis on bulk RNA-seq datasets that met our eligibility criteria (see Materials and Methods). Volcano plots (**Fig. 1a**) illustrate the significant DEGs for each primary dataset, with red and blue indicating up-regulation and down-regulation, respectively. While only a few DEGs overlapped between at least two datasets, and no common DEGs were found across all datasets, we identified numerous convergent BPs involved in the MDD pathogenesis (**Fig. 1b**). We identified 26 common BPs and 76 shared BPs among three datasets (Ramaker et al., 2017; Cathomas et al., 2022; Trang et al., 2018). These processes include cellular response to cytokine stimulus, second messenger-mediated signaling, sensory organ development, regulation of gene expression, and generation of neurons (**Fig. 1c**). Notably, these BPs are enriched by an interconnected network of immune- and nervous system-related DEGs derived from PBMCs and ACC samples (**Fig. 1d**).

**Fig. 1.**
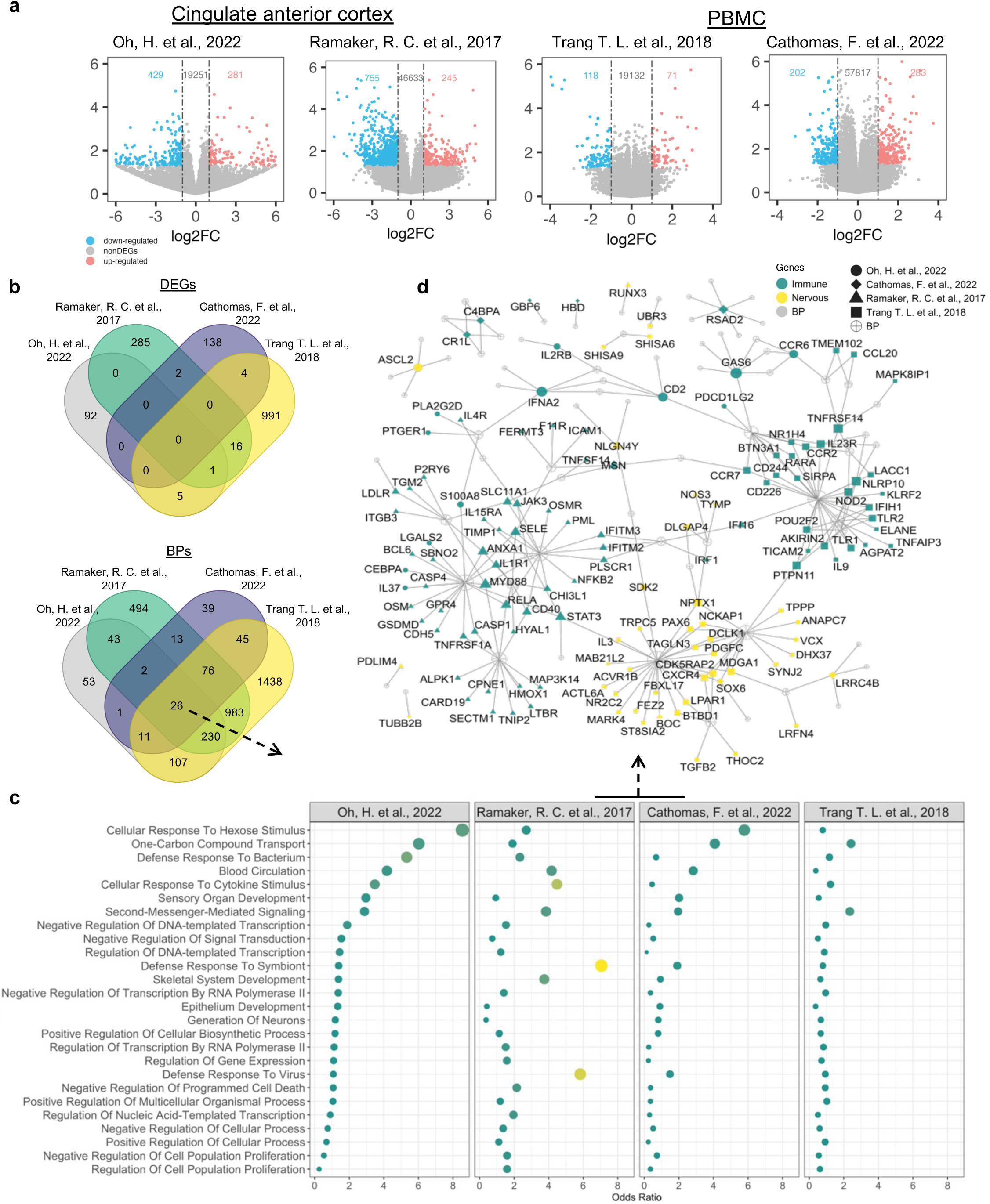
Characterization of common DEGs and biological processes across blood and brain (cingulate cortex) transcriptome datasets of MMD patients. (**a**) Volcano plots showing up- (red) and down-regulated (blue) differentially expressed genes (DEGs) across transcriptome of the following datasets: Oh, H. et al. (2022), Ramaker, R. C. et al. (2017), Trang, T. L. et al. (2018), and Cathomas, F. et al. (2022). (**b**) Venn diagrams illustrating the overlap of DEGs and biological processes (BPs), highlighting the commonalities among the studies. (**c**) Dot plots showing the 26 common BPs enriched by DEGs from each data set. (**d**) These latter are shown in a network view pointing to a neuroimmune interconnection throughout related BPs.

To further investigate the clustering of BPs enriched by DEGs from each study and to understand their inter-relationships, which could provide novel insights into the neuroimmunobiology of MDD, we performed gene enrichment analysis using the DEGs from each dataset individually. Visualization of the data through uniform manifold approximation and projection (UMAP) for dimensionality reduction revealed prominent clusters of neuroimmunological processes (**Fig. 2a**). Leveraging the rrvgo package to interpret lists of gene ontology (GO) terms, we gained an integrated perspective of various neuroimmunological BPs via a semantic similarity matrix (**Fig. 2b**). This correlogram suggests a complex interplay of neuroimmunological mechanisms underlying MDD.

**Fig. 2.**
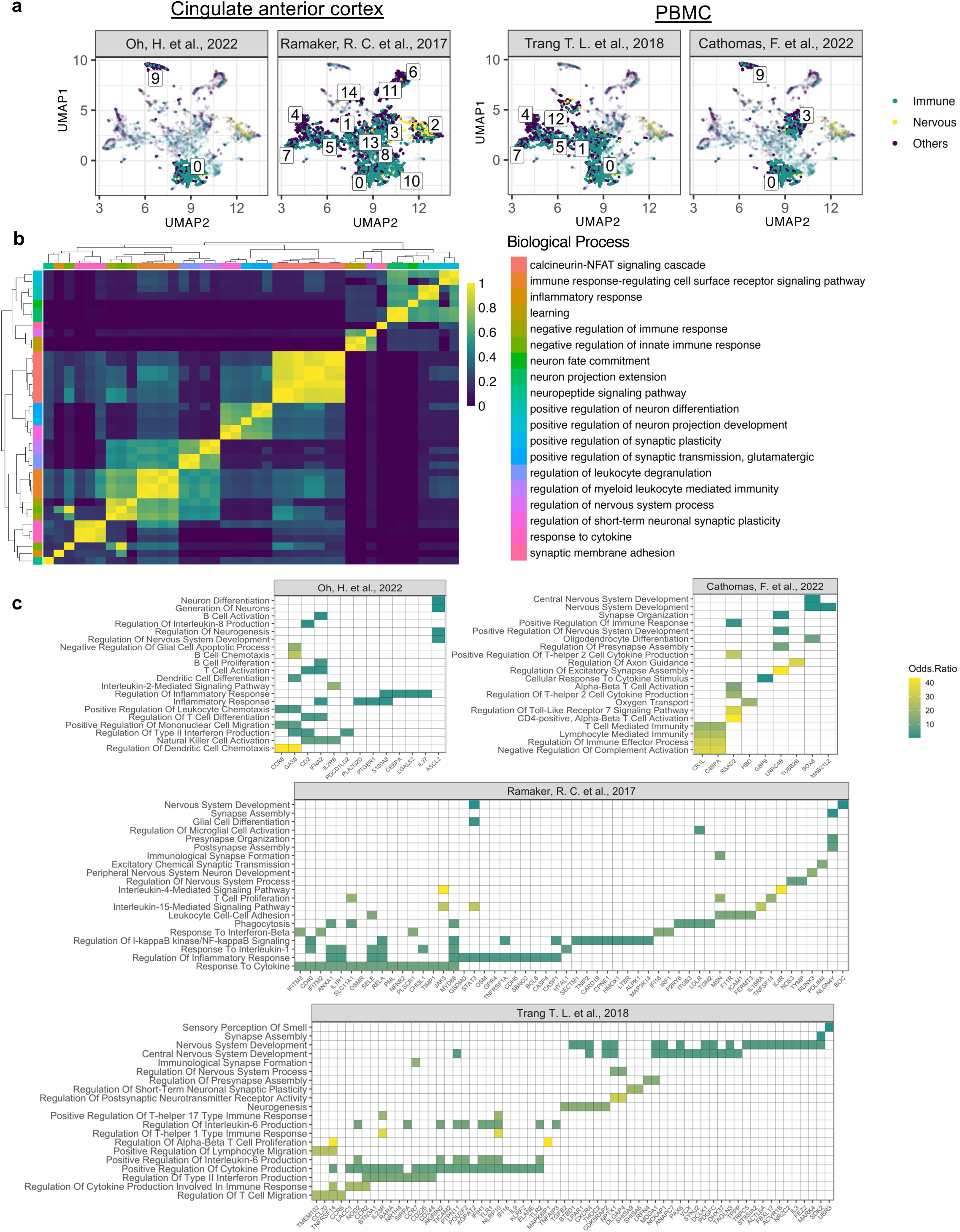
Clusters of neuroimmunological biological processes in blood and brain (cingulate cortex) transcriptome datasets of MMD patients. (**a**) Uniform Manifold Approximation and Projection (UMAP) exhibiting clusters of biological processes (BPs) enriched by DEGs from the following datasets: Oh, H. et al. (2022), Ramaker, R. C. et al. (2017), Trang, T. L. et al. (2018), and Cathomas, F. et al. (2022). Clusters enriched with neuroimmunological BPs are exhibited, as indicated by different numbers. (**b**) Correlogram illustrating a similarity matrix of neuroimmunological BPs enriched by DEGs from all MDD datasets. (**c**) Heatmaps showing DEGs associated with different neuroimmunological BPs found in each dataset.

Additionally, we performed gene annotation to explore the relationships between genes and BPs, assigning functional roles to each gene (**Figs. 2c** and **2d**). This analysis revealed that DEGs across the studies regulate key neurological BPs, including neuron generation and differentiation, nervous system development, synapse organization and assembly, sensory perception of smell, glial cell differentiation, oligodendrocyte differentiation, and the regulation of nervous system processes. The immune-related BPs involve DEGs in B and T cell activation, cytokine-related processes (e.g., IL-1, IL-2, IL-6, IL-8, IL-17, and interferons [IFNs]), as well as leukocyte chemotaxis and phagocytosis. Thus, while different genes may be dysregulated in various MDD cohorts, they frequently converge on similar biological pathways and processes, underscoring the complex and multifactorial nature of MDD.

### Overlapping genes associated with depression in genome-wide and transcriptome datasets

The findings of our consensus analysis indicate a systemic involvement of neuroimmunological processes across PBMCs and ACC, reinforcing the complexity and multifactorial nature of MDD. To enhance the robustness, accuracy, and generalizability of our findings, ultimately leading to a deeper and more reliable understanding of the biological basis of depression and highlighting potential targets for therapeutic intervention, we integrated findings from the most extensive worldwide GWAS meta-analysis Als et al., 2023^6^. This study included over 1.3 million individuals, with 371,184 diagnosed with depression. Furthermore, we also incorporated the most extensive worldwide transcriptomic meta-analysis findings by Wittenberg et al., 2020^8^, which included PBMC datasets of 1,754 MDD cases and 1,145 healthy controls (HC). Wittenberg and colleagues quantitatively reviewed 10 published whole-genome transcriptional datasets from PBMCs and whole peripheral blood leukocytes (PBLs) samples in case-control studies of MDD, focusing on studies developed before 2018.

The flow chart in **Fig. 3a** outlines the integrative approach taken to combine data from various sources. We performed a consensus analysis involving 411 genes significantly associated with depression (identified by Als et al., 2023)^6^. and transcriptome datasets from the ACC (Oh et al., 2022^13^, and Ramaker et al., 2017 ^14^), datasets of bulk RNAseq from PBMCs (Cathomas et al., 2022^15^, and Trang et al., 2018 ^16^), and the transcriptomic meta-analysis by Wittenberg et al., 2020)^8^, which found 343 genes with a false discovery rate <5%.

**Fig. 3.**
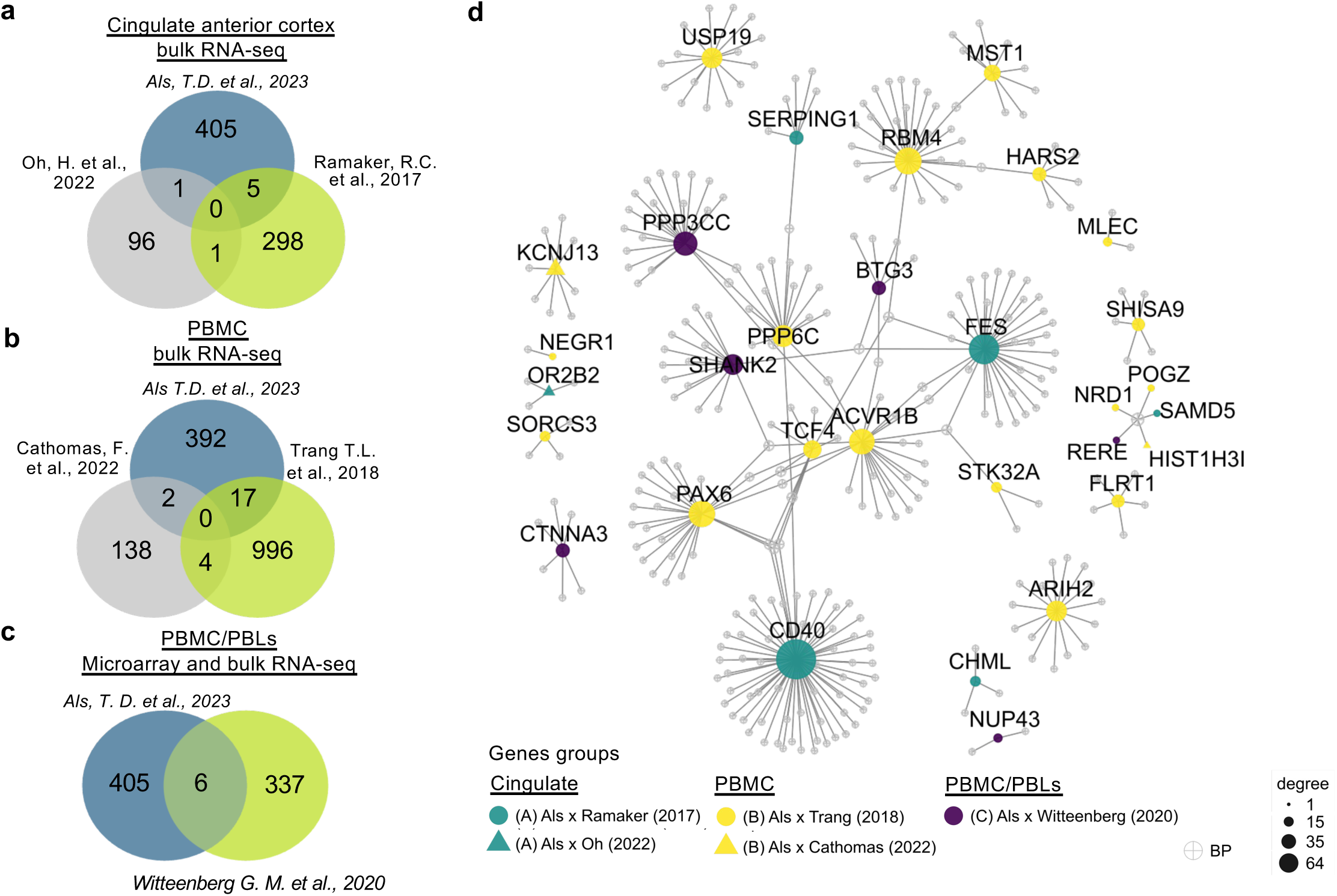
Overlapping genes associated with depression in genome-wide and transcriptome datasets. (**a**) Flow chart indicating the integrative systems analysis of primary data sources used for our study. From left to right: Als, T.D et al. (2023) (a; GWAS meta-analysis); the four transcriptome datasets that we analyzed here, two derived from brain cingulate cortex samples (Oh, H. et al. 2022, and Ramaker, R. C. et al. 2017), and two derived from blood samples (Tran, T. L. et al. 2018, and Cathomas, F. et al. 2022); and the Wittenberg et al. 2020 study providing data from a transcriptome meta-analysis of MDD patients. (**b-d**) The Venn diagrams exhibit the number of shared genes identified across each primary dataset. (**e**) The 31 shared genes are shown in the gene-biological processes (BPs) network. Yellow and green nodes represent differentially expressed genes from PBMCs and cingulate cortex datasets, respectively. Gray nodes represent BPs and edges the gene-BP association.

This approach identified 31 shared genes between genomic and transcriptomic studies (**Figs. 3b-d**), further visualized in a gene-BPs network (**Fig. 3e**), indicating their potential involvement in standard biological processes related to MDD. Six DEGs (Oh et al., 2022^13^, OR2B2; Ramaker et al., 2017 ^14^), CD40, CHML, FES, SAMD5, and SERPING1) were commonly identified in the GWAS meta-analysis and as DEGs in the ACC datasets (**Fig. 3b**). Additionally, 25 genes from the GWAS meta-analysis were identified as DEGs in the PBMC transcriptomes (**Figs. 3c, d**). Specifically, 17 genes overlapped with the Trang et al., 2018 ^16^ dataset (ACVR1B, ARIH2, FLRT1, HARS2, MLEC, MST1, NEGR1, NRD1, PAX6, POGZ, PPP6C, RBM4, SHISA9, SORCS3, STK32A, TCF4, and USP19), two genes with the Cathomas et al., 2022^15^ dataset (HIST1H3I and KCNJ13), and six DEGs (BTG3, CTNNA3, NUP43, PPP3CC, RERE, and SHANK2) were found in the Wittenberg et al., 2020^8^ transcriptomic meta-analysis. These results underscore the potential involvement of these shared genes in the molecular mechanisms underlying MDD and suggest specific gene relationships that may be critical in the disorder’s pathology.

### Altered neuroimmunological gene correlations underpin stratification of major depressive disorder

Next, we assessed the ability of neuroimmunological gene expression to distinguish MDD patients from HC, which could have implications for personalized medicine. Principal component analysis (PCA) indicated that these DEGs in the ACC (**Extended Data Fig. 2**) and PBMCs (**Figs. 4a, b**) stratify MDD patients from HC. Concentration ellipses and histograms visually represent the data distribution, indicating that the stratification was more pronounced when comparing DEGs from PBMCs than those in the ACC.

**Fig. 4.**
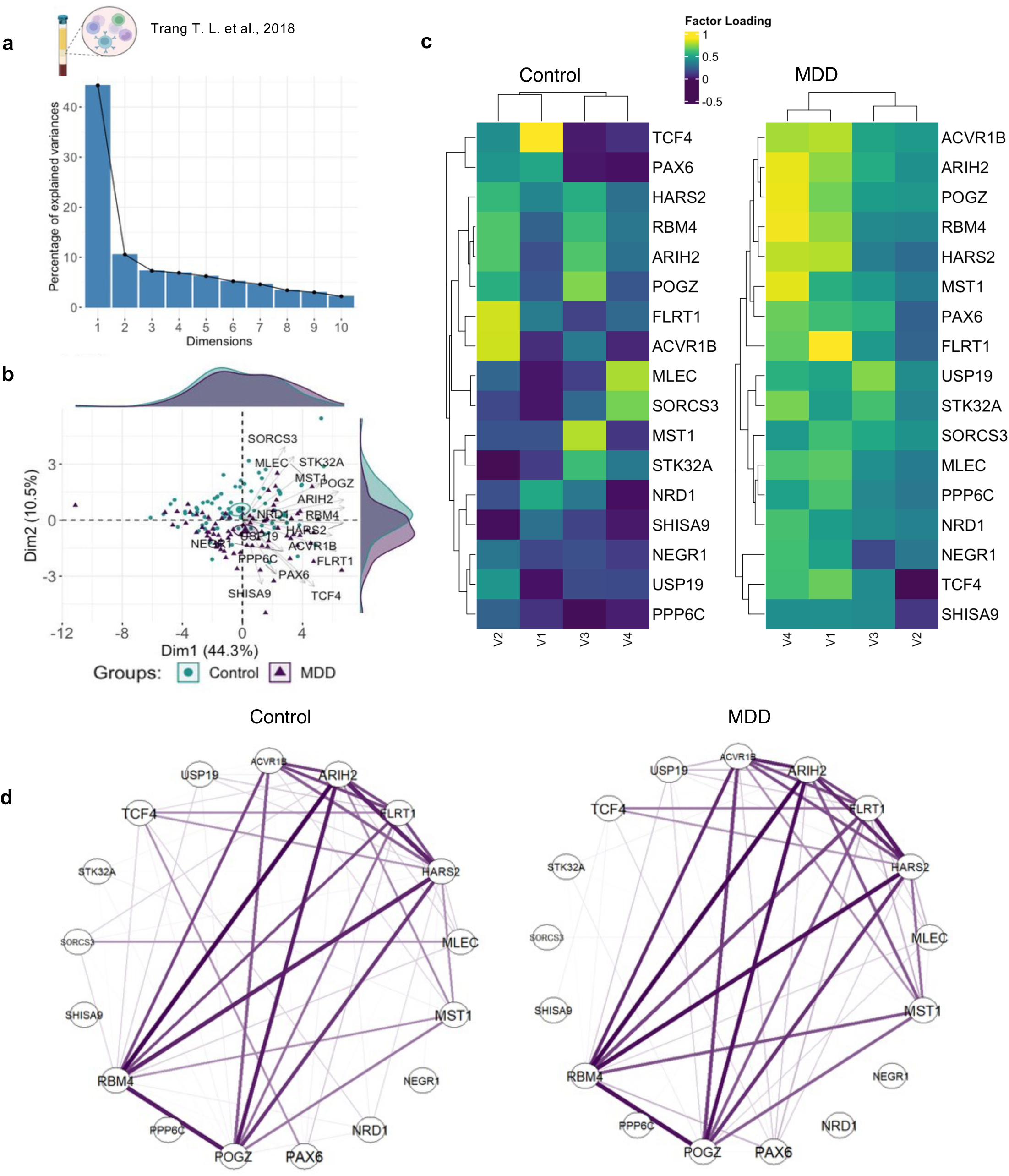
Stratification and relationships between shared genes across the genomic and transcriptomic primary MDD studies. (**a**) Bar plot displaying the percentage of explained variance for each principal component (PC) dimension from a Principal Component Analysis (PCA) with spectral decomposition. (**b**) PCA plot illustrating the stratification of MDD patients and healthy controls based on PBMCs. Genes with positive correlations align on the same side of the plot, while negatively correlated genes point in the opposite direction. Small circles represent concentration ellipses around the mean points of each group. The accompanying histograms depict the density distribution of samples (individuals) for each group. (**c**) Heatmaps obtained from exploratory factor analysis of the 17 shared genes with negative and positive loadings visualized for MDD patients and health controls from the Tang T. L. et al. dataset. (**d**) Correlograms show the topological correlation pattern among shared genes. The color scale bar represents the range of Spearman’s rank correlation coefficient for controls (left side of the graph) and MDD patients (right side).

Notably, the correlation distribution of variables (DEGs) in the PCA graphic highlighted correlated DEGs involved in stratifying MDD patients from HC. We conducted further analyses to elucidate the relationship patterns among the identified shared genes. This approach is crucial as it provides new insights into understanding the molecular underpinnings of MDD. For this purpose, we used the dataset from Trang et al., 2018 ^16^, which contains the highest number of shared DEGs with the GWAS meta-analysis. The Exploratory Factor Analysis (EFA) analysis revealed an overall increase in the factor loading of DEGs (**Fig. 4c**) in the MDD group.

Likewise, correlation analysis showed that while several correlations (based on Spearman’s rank correlation coefficient) were maintained, there were increased topological changes in correlation patterns among DEGs in MDD patients (**Fig. 4d**). For instance, there were increased connections between PAX6 and MST1 with other genes when comparing MDD to HC. However, we also observed a few reduced associations, including those between SORCS and MLEC genes. This detailed analysis underscores the complexity of the molecular interactions in MDD and highlights specific gene relationships that may play critical roles in the disorder’s pathology. Hence, these results indicate an interplay between DEGs in MDD through changes in systemic relationships.

### Probability of developing MDD based on gene expression

To elucidate biological aspects of DEGs in MDD from a systems biology perspective, we utilized a multivariate analysis of variance (MANOVA) to quantify the relative effects of gene expression on MDD concerning healthy controls. By leveraging the rich dataset from Trang et al., 2018 ^16^ the relative effect analysis pinpointed a subset of DEGs (e.g., SORCS3, PPP6C, NEGR1, and PAX6; **Fig. 5a**), which stood out due to the absence of overlapping confidence intervals in the analysis, indicating that they may play a more pronounced and distinct role in MDD. To ensure the robustness of our findings and to guard against the potential for spurious significance due to multiple comparisons, we conducted a rigorous false discovery rate (FDR) analysis. This rigorous statistical approach identified NEGR1, PAX6, PPP6C, and SORCS3 as DEGs in PBMCs, which remained significant after FDR correction (**Extended Data Fig. 3**). However, none of these five DEGs showed significant expression changes in ACC samples (**Extended Data Fig. 4**).

**Fig. 5.**
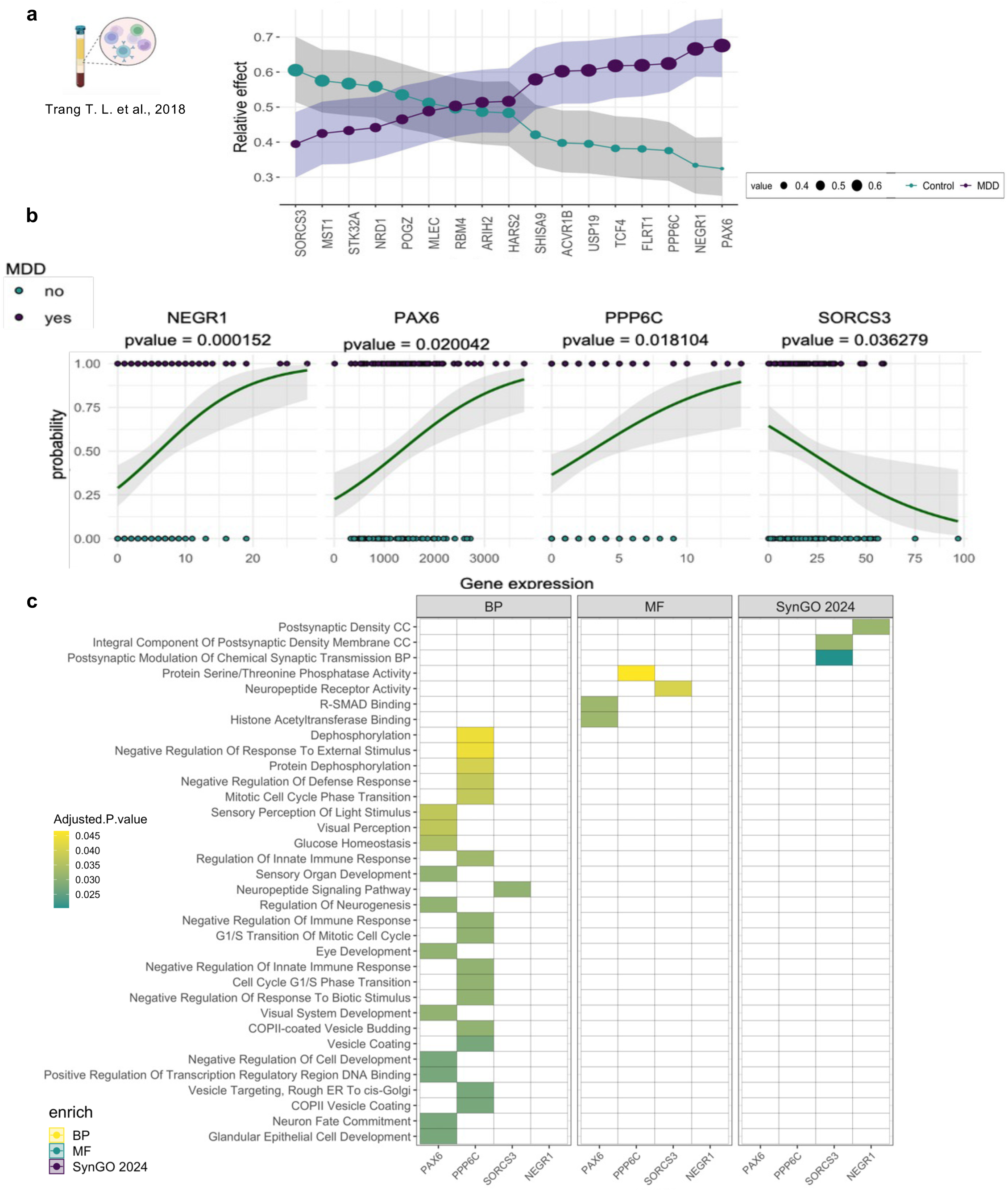
The probability of developing MDD based on the expression of NEGR1, PAX6, PPP6G, and SORCS3 genes. Relative effect of neuroimmunological genes in MDD. The graphic shows the relative effects (calculated using MANOVA test) of DEGs present in (**a)** PBMCs of MDD patients versus healthy controls. The circle size indicates the probabilistic measure (relative effect size). Shadows show confidence intervals. Turquoise and dark purple dots represent healthy individuals and MDD patients, respectively. (**b**) Scatter plots display the binomial logistic regression results for the expression of each gene *NEGR1*, *PAX6*, *PPP6G*, and *SORCS3*, across the MDD group compared to healthy controls (dataset: Trang T. L. et al., 2018). The MDD groups are represented with black color for "yes" (1 = presence of the MDD) and green color for "no" (0 = healthy controls). **(c)** The heat map exhibits the enriched biological processes (BP), molecular function (MF), and synapse-related processes based on the SynGO 2024 database for the four selected genes. The graphic at the bottom shows the number of enriched processes.

Having identified a set of DEGs that withstood the stringent FDR correction, we next sought to evaluate their predictive capacity in forecasting the risk of developing MDD. Through logistic regression analysis, we confirmed the potential significance of NEGR1, PAX6, PPP6G, and SORCS3 as predictive biomarkers for MDD (**Fig. 5b**). Simultaneously, the DEGs that did not survive the FDR correction did not demonstrate the same predictive power (**Extended Data Fig. 5**). Notably, the enrichment analysis of biological processes, molecular functions, and synapse-related processes associated with these four DEGs, as revealed by the SynGO 2024 database, not only reinforces their predictive value but also suggests a functional role in the neurobiological pathways in the PBMCs, to serve as valuable indicators of MDD risk (**Fig. 5c**).

Among the potential neuroimmunological roles ascribed to these four DEGs, NEGR1 and SORCS3 are implicated in various synaptic-related processes (**Fig. 5c**), as revealed by enrichment analysis of BPs, molecular function (MF), and synapse-related processes. Namely, SORCS3 is involved in neuropeptide receptor activity, while NEGR1 participates in synaptic organization and function. PPP6C, conversely, is associated with protein dephosphorylation, the negative regulation of defense response, and several cell-cycle-associated processes. PAX6, a transcription factor, plays a crucial role in neuron fate commitment, the positive regulation of transcription regulatory region DNA binding, the regulation of neurogenesis, glucose homeostasis, and visual perception. These diverse functional roles played by these DEGs suggest that they may be not only predictive biomarkers for MDD but also key molecules in the complex molecular networks that underlie the disorder’s pathophysiology.

### Diseasome of NEGR1, PAX6, PPP6G, and SORCS3

To gain a deeper understanding of the implications of NEGR1, PAX6, PPP6C, and SORCS3 in MDD, we conducted a comprehensive diseasome analysis. The diseasome conceptual framework maps the genetic and molecular connections between diseases, providing a systems view of how various disorders are interlinked through shared genetic factors. This approach revealed the gene-disease associations (GDAs) for the NEGR1, PAX6, PPP6C, and SORCS3 genes. PAX6 emerged as the gene most associated with different diseases, followed by NEGR1, SORCS3, and PPP6C (**Fig. 6a**). The heatmap (**Fig. 6b)** specifically highlights diseases related to at least two genes, revealing potential shared genetic factors among various mood and mental disorders and their comorbidities, such as intellectual disability, obesity, diabetes, and malignancies. Additionally, single nucleotide polymorphisms (SNPs) in these genes have been characterized and associated with MDD-related symptoms as reported in the GWAS catalog (**Figs 6c** and **6d**).

**Fig. 6.**
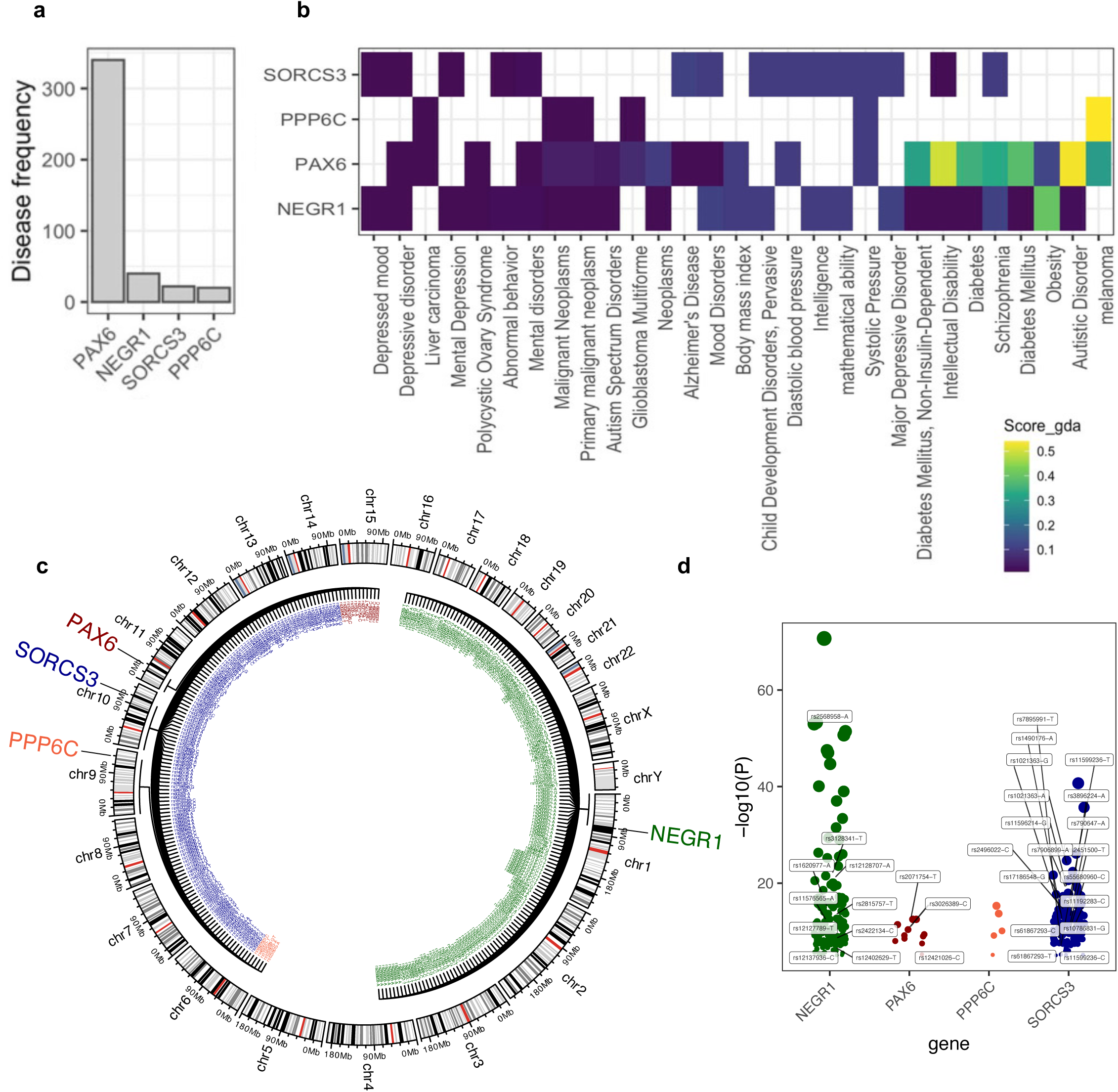
The diseasome of NEGR1, PAX6, PPP6G, and SORCS3 genes. (**a**) The bar plot exhibits the number of diseases associated with each gene. (**b**) The heatmap shows the gene-disease associations (GDAs) score for NEGR1, PAX6, PPP6G, and SORCS3 genes. The heatmap (**c**) shows diseases associated with at least two genes. (**c**) Circos plot depicting the chromosomal locations of the four genes: PAX6 (red), SORCS3 (blue), PPP6C (orange), and NEGR1 (green). SNPs associated with these genes, as reported in the GWAS catalog, are highlighted. (**d**) Dot plot representing the − log 10 p-values (y-axis) for all cataloged SNPs within PAX6, SORCS3, PPP6C, and NEGR1 (x-axis). The size of the dots is proportional to the statistical significance of the SNPs. (**d**) Dot plot illustrating the risk values associated with each SNP linked to the investigated trait.

### Consistent differential expression of PAX6 across blood and amygdala in MDD

To expand our understanding of neuroimmunological interactions between the blood and the central nervous system (CNS), we examined whether NEGR1, PAX6, PPP6C, and SORCS3 genes exhibit differential expression in CNS regions beyond the ACC. We aimed to identify common DEGs across these regions and compare them with the meta-analysis results of Als et al., 2023)^6^. Transcriptomic data for the amygdala, which is involved in emotional regulation, stress response, and mood processing, was obtained from the primary dataset of Labonté B. et al. (2017)^17^, while data for additional CNS regions were extracted from Li J.Z. et al. (2013)^18^.

We identified varying numbers of overlapping differentially expressed genes (DEGs) across different CNS regions, with the amygdala showing the highest number of DEGs (**Fig. 7a**). Notably, PAX6 was identified among the DEGs in the amygdala. In contrast, NEGR1, PPP6C, and SORCS3 did not exhibit significant differential expression in other CNS regions. This suggests that the expression of these genes may be region-specific within the CNS or that other factors, such as the local tissue environment or disease states, could influence their regulation. Nonetheless, several other DEGs identified in our study overlapped with those reported in the Als et al., 2023^6^ meta-analysis, highlighting critical genes implicated in both CNS and peripheral blood transcriptomes of MDD patients. For instance, genes such as PAX6, ACVR1B, POGZ, and FLRT1 were detected in the blood transcriptome dataset from Trang, T.L. et al. (2018). Additionally, KCNJ13 was identified as a DEG in Cathomas, F., et al. (2022), while CTNNA3 emerged as a DEG in the Wittenberg G.M et al., 2020^8^, (number) meta-analysis.

**Fig. 7.**
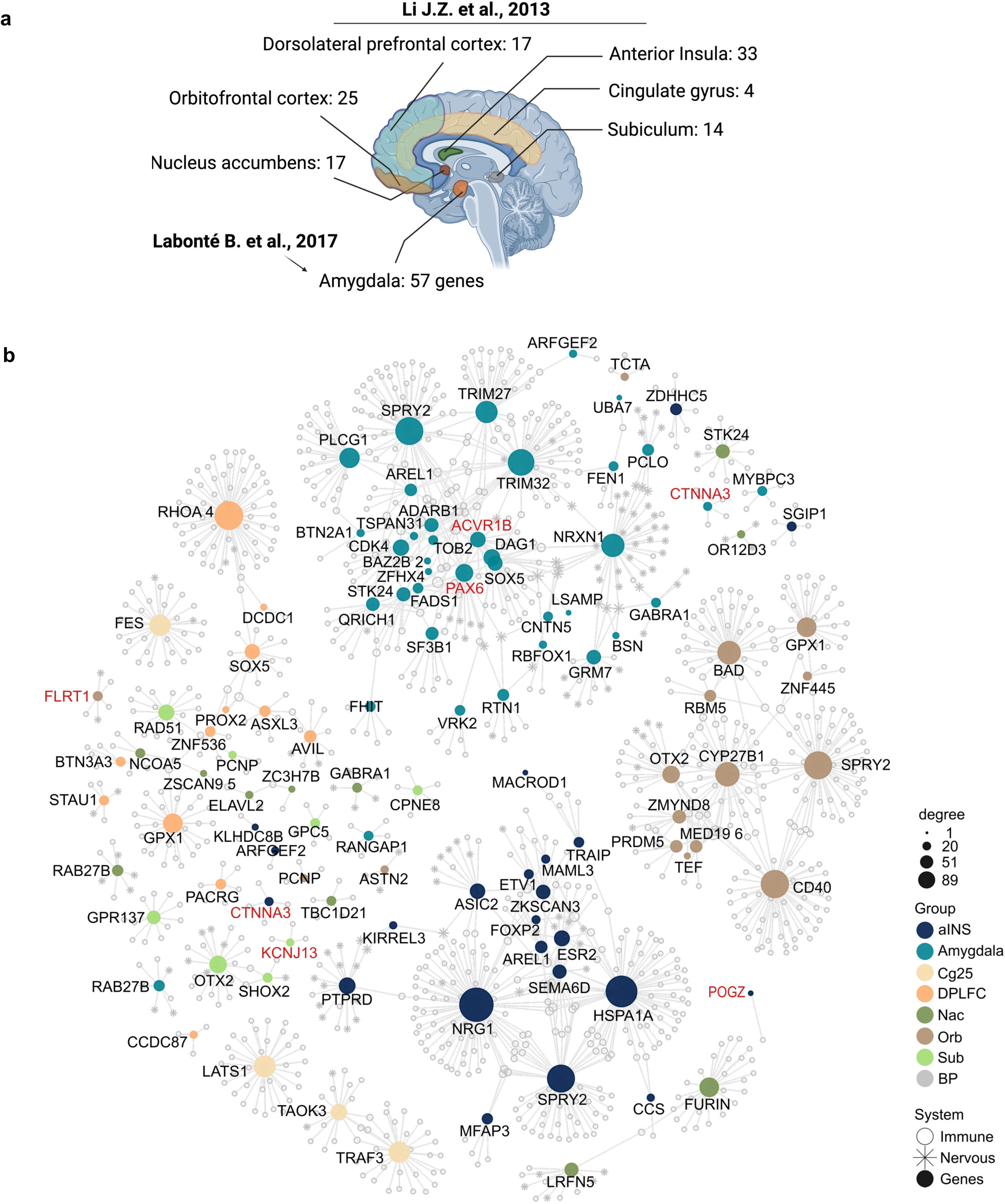
Differentially Expressed genes overlapping across various central nervous system regions with Als, T.D. et al., 2023 Meta-analysis. (**a**) The Fig. shows the names of different CNS regions along with the number of overlapping DEGs. Transcriptome data for the amygdala was obtained from the primary dataset by Labonté B. et al., 2017, while data for other CNS regions were sourced from Li J.Z. et al., 2013. (**b**) The network illustrates the DEGs overlapping with those identified in the Als, T.D. et al., 2023 meta-analysis. The PAX6, ACVR1B, POGZ, and FLRT1 genes are also present in the blood transcriptome data from Trang, T.L. et al., 2018. Additionally, KCNJ13 is identified as a DEG in the study by Cathomas, F. et al., 2022, and CTNNA3 is found as a DEG in the metanalysis by Wittenberg G.M. et al., 2020.

Taken together, the shared DEGs, particularly PAX6, point to potential biomarkers and therapeutic targets, providing a more integrative perspective on the systemic and neurobiological processes involved in MDD.

### Immunomodulatory effects of chronic restraint stress on PAX6 expression and immune cell populations in mice

The findings above suggested PAX6 as a critical predictor of MDD. Consequently, we extended our investigation by exploring stress-induced PAX6 expression in mice. To examine immune cell populations and PAX6 expression *in vivo*, we employed flow cytometric analysis of whole blood cells of mice exposed to chronic restraint stress (CRS), a well-characterized model for the induction of anhedonic behaviors in rodents^19^. Our analysis revealed four distinct immune cell clusters **(Fig. 8a-c)**, each delineated by the expression of specific markers (CD45, CD3, CD4, CD11b, CD19, Ly6C). Cluster 4 **(Fig. 8d-f)** exhibited an upregulation of PAX6 in immune cells expressing CD11b and Ly6C following CRS, suggesting a role for PAX6 in stress-mediated immune responses.

**Fig. 8.**
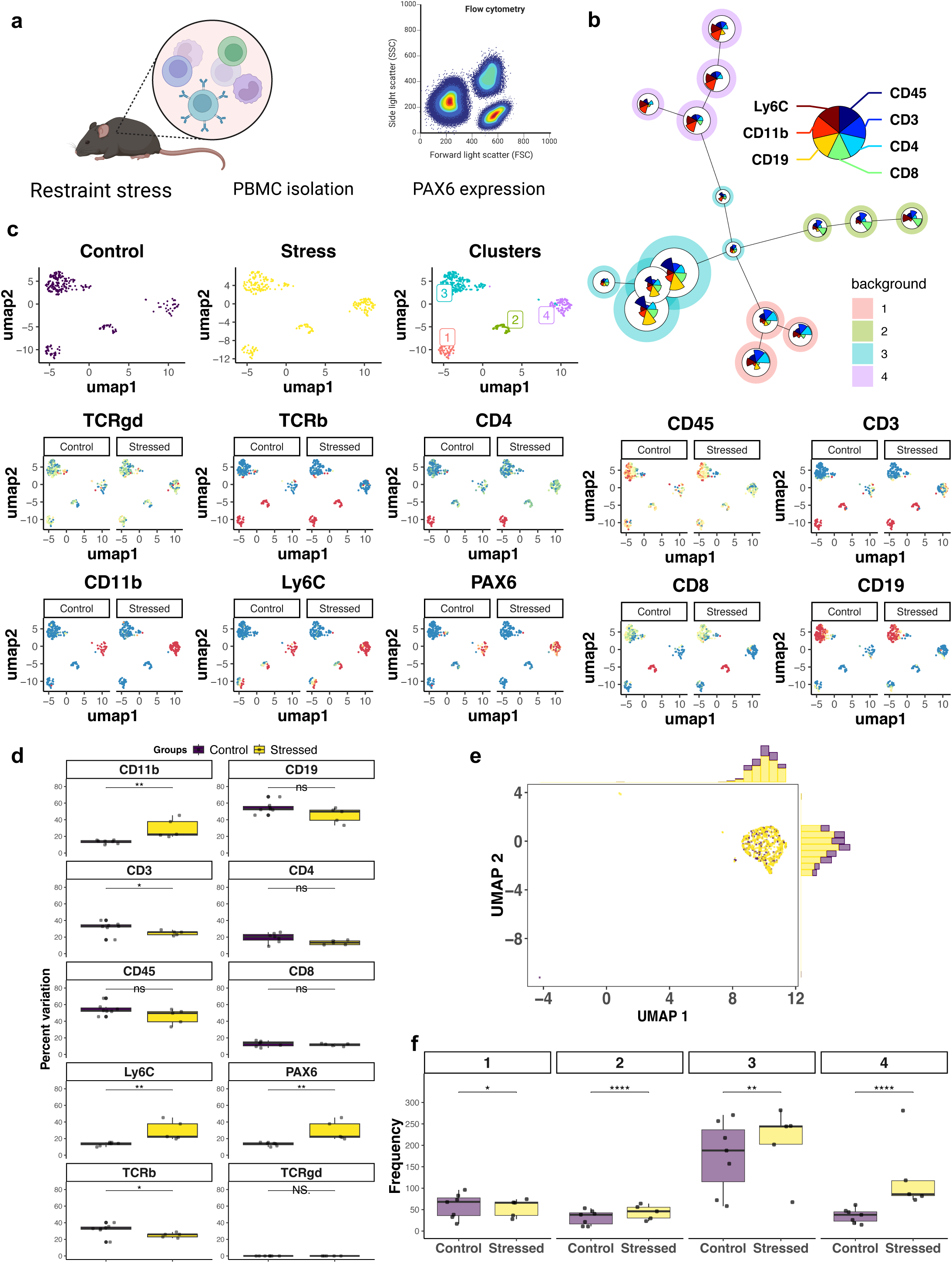
Immunophenotyping of Immune Cells in Male C57BL/6J Mice Subjected to Chronic Restraint Stress (CRS). (**a-c**) Flow cytometry analysis shows the identification of four distinct immune cell clusters based on the expression of CD45, CD3, CD4, CD11b, CD19, and Ly6C. (**c-f**) Detailed analysis of Cluster 4, highlighting the upregulation of PAX6 expression within immune cells following CRS. The Fig. illustrates the immunomodulatory effects of CRS, with an increase in PAX6 expression and the expansion of myeloid and plasma cell populations (CD11b and Ly6C), as well as a reduction in T cell markers (CD3). N=7 controls and 5 stressed. For (**f**), student’s t-test was used. *p<0.05, **p<0.01, ****p<0.0001.

The upregulation of PAX6 in CD11b and Ly6C-expressing cells was accompanied by an increase in the frequency of cells CD11b+Ly6C+ cells, indicating an expansion of inflammatory myeloid cells (monocytes and neutrophils) and plasma cells. Conversely, we observed a reduction in CD3 and TCRα/β expression, which are T cell markers, indicating a potential suppression of cellular adaptive immune response due to stress. These findings collectively suggest that stress can modulate the immune system through various mechanisms, influencing both innate and adaptive immune responses.

## DISCUSSION

To our knowledge, this work represents the first dataset-based work integrating genomics, transcriptomics, and immunophenotyping validation of leukocyte-expressing PAX6 as a critical predictor of MDD. A key strength of our study is the enhanced robustness and generalizability achieved through data integration of the most extensive worldwide-published GWAS meta-analysis^6^, encompassing over 1.3 million individuals, and the most comprehensive PBMC transcriptomic meta-analysis by Wittenberg et al.^8^.

Broadly, our findings align with a recent integrative analysis of transcriptional datasets using the RobustRankAggreg package (RRA). Zhong et al.^24^ investigated the correlation between immune-associated DEGs in PBMC and ACC datasets. They also found more robust DEGs in the blood than ACC. However, as described here, the 25 overlapping genes found in PBMCs form a network of classical molecules associated with the nervous system. These DEGs stratify between MDD and healthy control groups in PCA, with a more pronounced separation between groups using PBMC-derived DEGs than those from the ACC.

To gain new insights into the MDD pathophysiology, correlation analysis identified altered connections between the common genomic and transcriptomic DEGs in PBMCs. For instance, the increased connections of PAX6 and MST1 with other genes may implicate these as central hubs in MDD pathology, potentially influencing multiple downstream biological processes. PAX6, known for its role in neural development^25^, and MST1, involved in apoptosis and immune response^26^, indicate new neuroimmunological pathways that could be critical in the development and progression of MDD. Similarly, the reduced associations between SORCS3, which is involved in neuronal development and synaptic function^27,28^, and MLEC, an M1 to M2 macrophage polarization promoter^29^, suggest potential dysregulation or shifts in molecular interactions that merit further investigation.

These findings suggest a dynamic relationship between the immune and nervous systems that extends beyond the known "crosstalk" between both systems involving the release and diffusion of neurotransmitters from nervous tissue to regulate immune cells through leukocyte-surface receptors^30–33^. Consistent with this, immune cells can also synthesize and release neurotransmitters^34,35^, including acetylcholine (ACh)^36–38^, serotonin^39–41^, and glutamate^42^ that modulate cell activation, acting as autocrine or paracrine modulators.

Growing evidence supports the association of NEGR1, PAX6, PPP6G, and SORCS3 genes with MDD. For example, previous research has implicated PAX6 in neurodevelopmental processes and visual perception^43,44^, which are relevant to the cognitive and sensory symptoms observed in MDD^45,46^. Moreover, the involvement of PPP6C in protein dephosphorylation and cell-cycle processes^47^ aligns with existing literature suggesting dysregulation of cell-cycle signaling^48^ in MDD. Additionally, the involvement of NEGR1 and SORCS3 in synaptic function and neuropeptide receptor activity^27,28,49^, alongside their associations with other mental disorders^50^, suggests that synaptic dysregulation could be a shared mechanism.

Utilizing a diseasome analysis, we mapped the GDAs for NEGR1, PAX6, PPP6C, and SORCS3, revealing their potential roles in a spectrum of human disorders. This approach provides a valuable systems-level perspective on MDD pathophysiology. The shared genetic associations of NEGR1, PAX6, PPP6C, and SORCS3 with other diseases underscore the complex interplay between genetic factors in MDD and other conditions.

The association of PAX6 with intellectual disability^43^, for instance, suggests that disruptions in neuronal development and differentiation might be a common pathological feature. Similarly, NEGR1’s linkage to obesity and diabetes indicates^51,52^ a broader role in metabolic processes that could intersect with mood regulation pathways. PPP6C, associated with protein dephosphorylation and immune regulation^47^, links to malignancies, highlighting a potential intersection between cellular signaling, immune responses, and mood disorders.

This overlap emphasizes the need for a holistic approach to understanding MDD, considering its unique and shared pathophysiological aspects. It suggests that MDD may share genetic underpinnings with other disorders, such as obesity, diabetes, and malignancies, suggesting potential links between MDD and these seemingly disparate conditions, which could explain the high rates of comorbidity observed clinically^53^,

Consistent with the interconnection between the nervous and immune systems, our findings in a mouse model of CRS suggested that in addition to its role in neurogenesis, PAX6 may also be implicated in the immune response, particularly under conditions of psychological stress. The observed increase in PAX6 expression, alongside heightened levels of Ly6C and CD11b and a reduction in CD3 lymphocytes, indicate that PAX6 could be a potential biomarker for depression, specifically linking the neurogenic and immune systems. The increase in Ly6C and CD11b expression suggests a shift toward a pro-inflammatory state, while the reduction in CD3 lymphocytes points to a potential suppression of adaptive immunity. This novel association opens new avenues for research into the role of PAX6 in immune cells, an area previously unexplored.

Given these findings, we hypothesize that PAX6 may be involved in a novel biological process within immune cells, potentially regulating the balance between innate and adaptive immunity under stress conditions. This uncharted role of PAX6 warrants further investigation, as it could reveal new molecular targets for therapeutic interventions in stress-related disorders like depression. Future research should focus on elucidating the specific pathways by which PAX6 influences immune cell function and how these changes correlate with the onset and progression of depression. Understanding the dual role of PAX6 in both neurogenesis and immune regulation could significantly advance our knowledge of the mind-body connection and the development of new treatment strategies for mental health disorders.

In conclusion, we not only identified a set of DEGs with pronounced effects on MDD risk and potential as biomarkers, but also delved deeper to understand their systemic implications within the context of disease networks. Understanding how these genes contribute to the neuroimmunological processes in MDD could unveil novel insights into the disorder’s etiology. By elucidating the molecular underpinnings of MDD, we can move closer to developing personalized treatment strategies that address the specific biological mechanisms underlying each patient’s condition.

Future studies are warranted to functionally validate the roles of these genes and their interactions in MDD pathogenesis. Additionally, future research should focus on validating these findings in larger, independent cohorts and exploring the functional implications of these altered gene relationships. This knowledge can pave the way for developing novel diagnostic tools and therapeutic strategies targeting these altered molecular pathways.

## ONLINE METHODS

Extended Data information, including demographic characteristics of MDD patients and the input/output transcriptomic data used to generate the Figures using R programming language, is available in Supplementary Tables.

### Data Curation

#### Genomic Dataset

To identify genetic changes associated with MDD, we used the genes associated with depression across the genome, as recently published by Als et al., (2023)^6^, as a reference. This association was determined using the Multi-Marker Analysis of GenoMic Annotation (MAGMA) tool, as implemented in the Functional Mapping and Annotation of Genome-Wide Association Studies (FUMA) platform v.1.3.6a^54,55^. This approach characterized 411 genes significantly associated with depression after Bonferroni correction for the number of genes tested.

#### Transcriptomic datasets

We searched for eligible transcriptome datasets published in the online library PubMed (https://pubmed.ncbi.nlm.nih.gov/) and Gene Expression Omnibus (GEO)^56^ indexed under the terms "major depressive disorder" and "human." We found 30 studies reporting whole-genome transcriptional data.

Studies were subsequently excluded based on the following criteria: 1) studies available only in original articles and not in databases (n = 6), 2) studies using snRNA-Seq (single-nucleus RNA sequencing) (n = 1), 3) studies addressing other categories of depression besides MDD (n = 1), 4) datasets from neurons, astrocytes, and immortalized B lymphocyte cell lines (n = 11), and 5) datasets with fewer than 50 significant differentially expressed genes (DEGs) (n = 1). This resulted in 10 eligible datasets published in 10 different articles, which were accessible for reanalysis. Two of these transcriptome datasets were derived from the ACC of patients with depression and healthy controls (Oh, H. et al. [2022] - GSE193417^13^ and Ramaker, R. C. et al. [2017] - GSE80655^14^) from bulk RNA-seq, a dataset of 6 brain regions including Anterior Insula (aINS), Cingulate Gyrus 25 (Cg25), Dorsolateral Prefrontal Cortex (DLPFC), Nucleus Accumbens (Nac), Orbitofrontal Cortex (OFC), Subiculum (Sub) of patients with depression and healthy controls (Labonté et al. [2017] - GSE102556) from bulk RNA-seq^17^, another from microarray including the amygdala of patients with depression and healthy controls (Li J. Z. et al. [2013] - GSE45642)^18^, while other two datasets obtained transcriptome data of PBMCs from MDD patients and healthy controls (Cathomas, F. et al. [2022]^15^ - GSE18855 and Trang T. L. et al. [2018]^16^ – (data available at GitHub: https://github.com/insilico/DepressionGeneModules) from bulk RNA-seq.

We also incorporated the meta-significant genes identified by Wittenberg G. M. et al. (2020)^8^, who evaluated four datasets from different sequencing methods, to enhance the robustness of our analyses using bulk RNA-seq and microarray data.

### Differential expression and functional enrichment analyses

The read counts of each dataset were transformed into log2 counts per million (CPM), and DEGs between the groups (MDD patients versus healthy controls) were identified using the DESeq2 pipeline in R (v. 4.3.2)^57^, with an adjusted *p*-value < 0.05, ensuring that only the most robust and statistically significant expression changes were considered in the analysis.

Functional enrichment analysis of biological processes (BPs) was performed using gene ontology analysis. For this purpose, we utilized the 31 genes common to both genomic and transcriptomic data associated with depression. BPs were identified using the clusterProfiler package^58^ in R programming. Enriched BPs were defined as significant based on an adjusted p-value < 0.05. The significance and magnitude of biological term enrichment are visualized in dot plots graphs generated with clusterProfiler. Furthermore, we utilized network graphs to illustrate interactions between enriched terms and DEGs, employing the ggnet2 package^59^ in R (described below). This approach provides a comprehensive view of complex biological relationships, enhancing our understanding of the biological processes underlying depression.

To visualize neuroimmune interactions, we filtered significant gene enrichment associations (p-value < 0.05) and categorized enriched BP into "neuro," "immune," or "other." Subsequently, we created a network object to display interactions (edges) between BPs and genes (nodes) using ggnet2^59^ in R.

### Analysis of neuroimmune clusters

To evaluate the neuroimmune clusters formed by BPs, we utilized the web tool "appyters," integrated with EnrichR^60^. This approach enables the identification of enriched BPs. It organizes them into clusters on a scatter plot, facilitating the visualization of their interactions with biological processes and among themselves, using the Leiden algorithm for clustering, resulting in a reduction in Uniform Manifold Approximation and Projection (UMAP) dimensional.

### Principal Component Analysis

Principal Component Analysis (PCA)^61^ with spectral decomposition^62^ was used to assess the ability of transcripts to differentiate between patients with MDD and healthy controls. This analysis was performed employing the prcomp and princomp functions from the factoextra package^63–65^ using R. These tools facilitate PCA by decomposing the dataset into principal components, aiding in visualizing and understanding the variance within the data, thus evaluating the effectiveness of the genes in distinguishing between the different groups.

### Correlation Analysis among DEGs

We utilized the multivariate analysis of correlations among DEGs in each group to identify and visualize correlations between different DEGs within each group (controls and MDD), as previously described^66^. The corrgram package^67,68^ in R created correlation plots, clearly representing the relationships among the DEGs. The psych package^69^ was employed for calculating correlations and conducting additional statistical analyses, while inlmisc^70^ facilitated data manipulation and presentation. Additionally, correlations among DEGs were visualized in circular network formats using the qgraph package^71^ in R.

### Exploratory Factor Analysis (EFA)

A critical aspect of Exploratory Factor Analysis (EFA) is uncovering underlying structures in real-world problems^72^. Specifically, R-mode methods of EFA aim to investigate the relationship between variables. The Dandelion Plot is an innovative method for visualizing EFA in R^72^, providing a more adequate representation of factors.

### Relative effect of gene expression on MDD

The relative effect of NEGR1, PAX6, PPP6C, and SORCS3 gene expression on the MDD phenotype was evaluated using Multivariate Analysis of Variance (MANOVA). Statistical analysis was conducted using the R packages npmv^73^ and reshape^74^, as previously described^75^. This approach allows for determining if there are statistically significant differences between groups concerning dependent variables. It is instrumental when multiple dependent variables are suspected to be correlated. Additionally, we computed confidence intervals for the relative effects using bootstrap techniques involving 1000 resamples. The data obtained were visualized using ggplot2^76^.

### False Discovery Rate Analysis

False Discovery Rate (FDR) analysis^77^ compared gene expression between different diagnoses using the packages dplyr^78^, ggplot2^76^, rstatix^79^, and ggpubr^80^. Initially, we grouped the data by genes and performed t-tests to compare expression means between diagnoses, adjusting p-values using the FDR method. The results were visualized using boxplots, highlighting significant differences in expression between diagnoses according to adjusted p-values < 0,05, generated usinf the ggplot^76^ function in R.

### Binomial logistic regression

We employed binomial logistic regression^81^ to analyze the relationship between the expression of the four genes with predictor variables with FDR < 0.05 (NEGR1, PAX6, PPP6C, and SORCS3) and the binary outcome of MDD as a control condition. The glm function was used from the base R package for logistic regression. Statistical significance was determined using a p-value < 0.05.

### NEGR1, PAX6, PPP6C, and SORCS3 association with MDD and diseasome analyses

To further evaluate the association of NEGR1, PAX6, PPP6C, and SORCS3 with the MDD phenotype, we utilized data from the GWAS Catalog. We filtered the SNPs related to these genes based on their p-values and risk scores. The genomic locations of the genes and their associated SNPs were represented using a circos plot generated with the R circlize package^82^. Additionally, we visualized the significance, risk, and SNP characteristics using R ggplot2^83^ plots to provide detailed insights into their roles in MDD.

Moreover, we assessed the association of NEGR1, PAX6, PPP6C, and SORCS3 with MDD and comorbidities by evaluating the diseasome of these four genes using Dysgenet^84^ through EnrichR^60^. This approach allows the integration of information on genes and their associations with various genetic diseases, disorders, and syndromes, compiling data from scientific literature and genetic variant databases. The creation and visualization of the diseasome network were conducted using the "ggnet2"^85^ package in R, which correlates genes with diseases, disorders, and syndromes, where nodes represent genes and edges represent relationships between them.

### Stress-induced expression of PAX6

#### Mice

All animal experiments were approved by the BWH IACUC. Eight-week-old male C57BL/6J mice were purchased from the Jackson Laboratory (Strain #000664) and housed in a conventional specific pathogen-free facility at the Hale Building for Transformative Medicine, Brigham and Women’s Hospital, Harvard Medical School. Mice were group-housed under a standard light cycle (12h light/dark) at 20-23C and humidity (∼50%) with *ad libitum* access to water and food.

#### Chronic restraint stress (CRS) model

This method is widely utilized to induce psychological stress in rodents. It is frequently employed in studies examining the effects of stress on behavior, physiology, and molecular biology, including models of depression and anxiety^86^. Tubes were washed with 70% ethanol before every use. 30-40 air holes were drilled using a 1/16” drill bit into 50-mL conical tubes (Falcon). Mice were placed into tubes and placed horizontally into a tube rack for 6 hours each day for 7 consecutive days. Stress sessions were started daily between 9-10 AM. After restraint, mice were returned to their cages, and tubes were washed with soap, water, and 70% ethanol.

#### Flow cytometric analysis

Blood was removed from the aorta with a 21G needle (Becton Dickinson, #305165) and placed in a blood collection tube containing the anti-coagulant heparin (Becton Dickinson, #365965) on ice. Briefly, blood was then resuspended in 1 mL of ACK lysing buffer (Thermo Fisher Scientific) for 5 min. Cells were centrifuged at 1600 rpm for 5 min, and supernatants were discarded. Cells were then stained for flow cytometric analyses using the following dyes/antibodies: fixable viability dye Zombie Aqua (1:1000; BioLegend; cat.# 423101) to exclude dead cells, BUV395-anti-CD45 (BD Biosciences; 1:200; cat.# 564616), BUV737-anti-CD3 (BD Biosciences; 1:200; cat.# 612771), PE-Cy7-anti-TCRb (Biolegend; 1:200; cat.# 109222), BV421-anti-TCRgd (Biolegend; 1:200; cat.# 118120), BUV496-anti-CD4 (BD Biosciences; 1:300; cat.# 612952), BV711-anti-CD8 (Biolegend; 1:300; cat.# 100748), AF700-anti-CD19 (Thermo Fisher Scientific; 1:300; cat.# 56-0193-82), APC-anti-CD11b (Biolegend; 1:100; cat.# 101212), BV785-anti-Ly6C (Biolegend; 1:100; cat.# 128041) and AF488-anti-Pax6 (Bioss USA; 1:100; cat.# bs-11204R). Surface markers were stained for 25 min at 4°C in Mg^2+^ and Ca^2+^ free HBSS with 2% FCS, 0.4% EDTA (0.5 M), and 2.5% HEPES (1M), then were fixed in Cytoperm/Cytofix (eBioscience), permeabilized with Perm/Wash Buffer (Thermo Fisher Scientific). The flow-cytometric acquisition was performed on an AURORA (Cytek) instrument by using DIVA software (BD Biosciences), and data were analyzed with FlowJo software version 10.1 (TreeStar Inc).

Data from 31,000 cells per mouse were processed using R 4.4.0 and RStudio 2024.04.2. The raw data underwent rigorous pre-processing, including compensation with the flowCore library, standardized logical transformation (w = 0.5, t = 1,000,000, m = 4.5), and z-score normalization to ensure consistency. Cells were then concatenated based on a 10-color panel: CD3, CD4, CD8, TCRγδ, TCRβ, CD11b, CD19, Ly6C, CD45, and PAX6, categorized into Control and Stressed groups.

Differential analysis was performed using FlowSOM, focusing on lineage-defining markers (CD19, CD3, CD4, CD8, CD11b), identifying four meta clusters. Initial cluster delineation was guided by manual gating in Kaluza Software with Radar plots (Fig. S1). UMAP provided a dimensional visualization of marker density and population distribution across groups and clusters.

## Author Contributions

HDD, ASA, ALN, IAFB, DLMF, LFS, IFS, PM, FYNV performed data and bioinformatics analyses; MGO and ENC performed CRS experiments; GCM, HN, RB, and CH provided scientific insights; HDD, CH, MAW, RMR, and OCM designed the study, wrote the manuscript and supervised the project.

## Acknowledgments

We thank the São Paulo Research Foundation (FAPESP grants 2018/18886-9 to OCM, 2020/16246-2 and 2023/13356-0 to DLMF; 2023/12268-0 to ASA, 2023/07806-2 to ISF, 2023/06086-6 to PM) for financial support. We acknowledge the National Council for Scientific and Technological Development (CNPq) Brazil (grants: 309482/2022-4 to OCM and 102430/2022-5 to LFS). MAW acknowledges support from the NIMH, NIDA, and NINDS (R01MH130458, R01MH132632, R01DA061199, R00NS114111), the Boston Claude D. Pepper Older Americans Independence Center, and the Brigham Research Institute. We thank Coordination for the Improvement of Higher Education Personnel (CAPES) (88887.699840/2022-00 to FYNV, 88887.801068/2023-00 ALN, 88887.992612-2024-00 IAFB) for financial support.

