## Supplementary material for "Integrative systems neuroimmunology reveals leukocyte-expressing PAX6 as a critical predictor of major depressive disorder": Supplemetary Figures

**Suppl. figure 1**

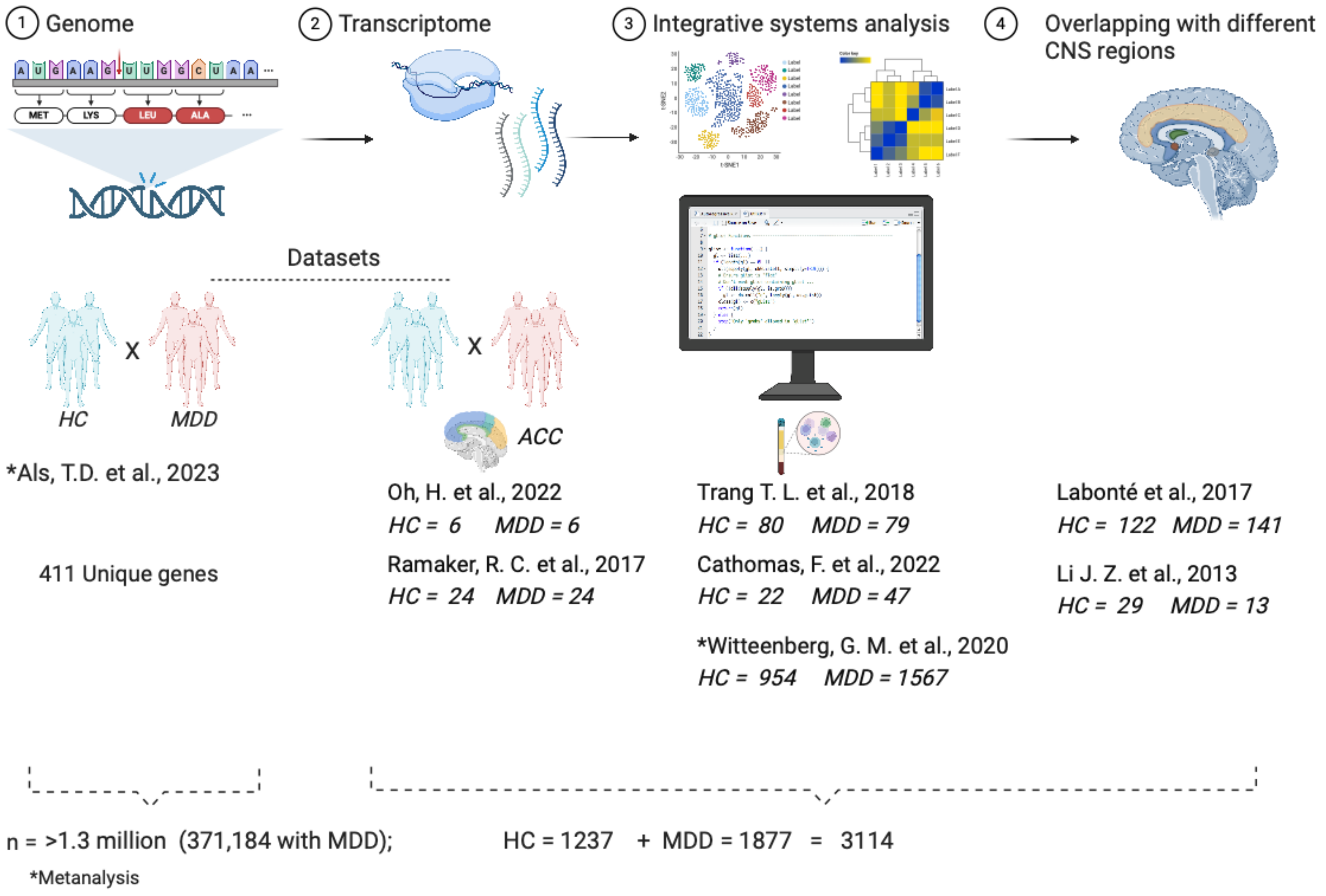


**Supplementary figure 1. Overview of primary data sets and study workflow.** Flow chart illustrating the integrative systems analysis of primary data sources utilized in our study. From left to right: Als et al. (2023), a GWAS meta-analysis identifying MDD-associated genes; these findings were compared with transcriptomic data, including blood and ACC transcriptomes, the Wittenberg et al. (2020) study provided transcriptome meta-analysis data from MDD patients, alongside datasets from various central nervous system regions.

Ramaker et al., 2017

**Suppl. figure 2**

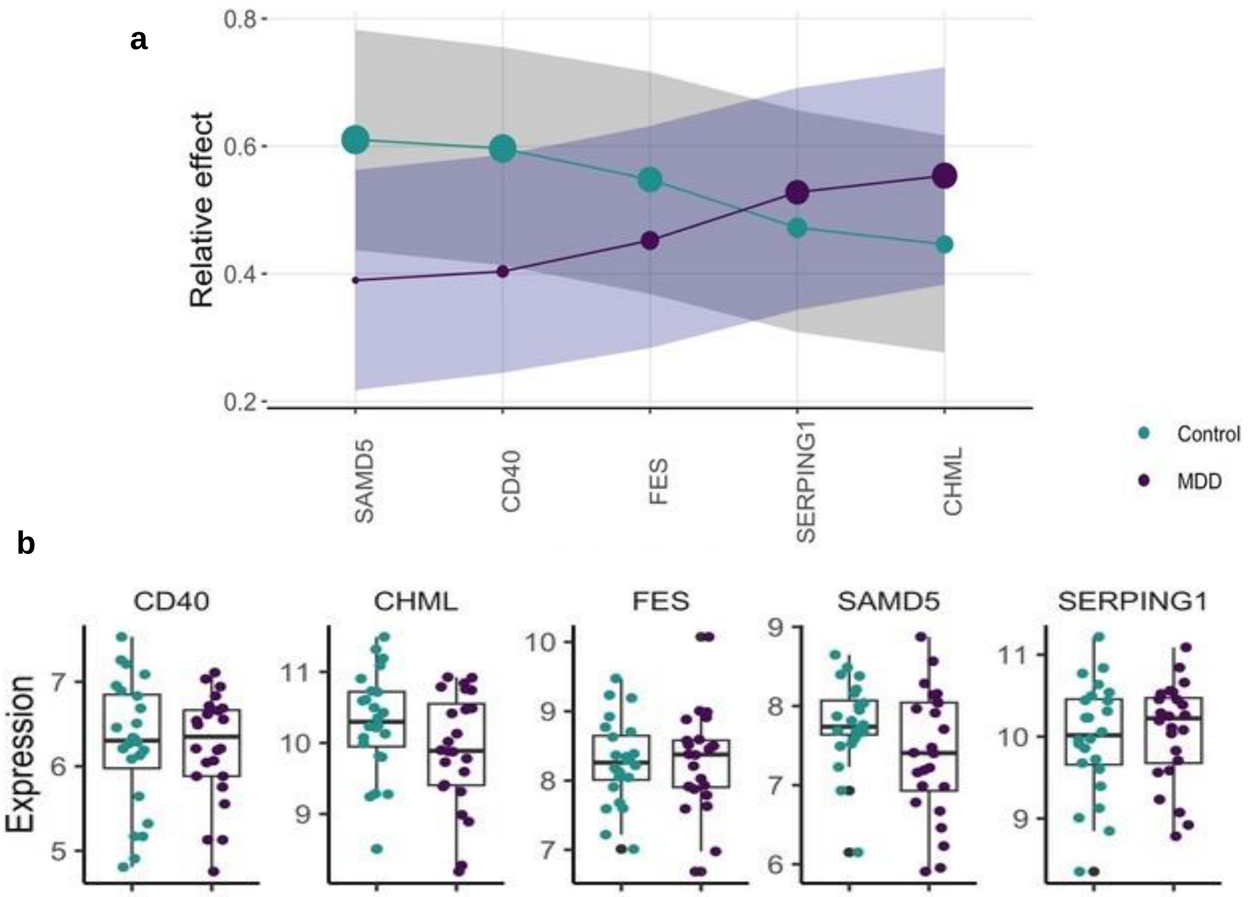


**Suppl. Figure 2. Relative effect of neuroimmunological genes in ACC of MDD patients.** The graphic shows the relative effects (calculated using MANOVA test) of DEGs present in (**a**) cingulate anterior cortex of MDD patients versus healthy controls. The circle size indicates the probabilistic measure (relative effect size). Shadows show confidence intervals. (**b**) Box plots of DEGs present in CAC of MDD patients compared to healthy controls after false discovery rate (FDR) correction for multiple comparisons. The analyses were performed with the primary dataset from Ramaker et al., 2017.

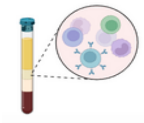

Trang T. L. et al., 2018

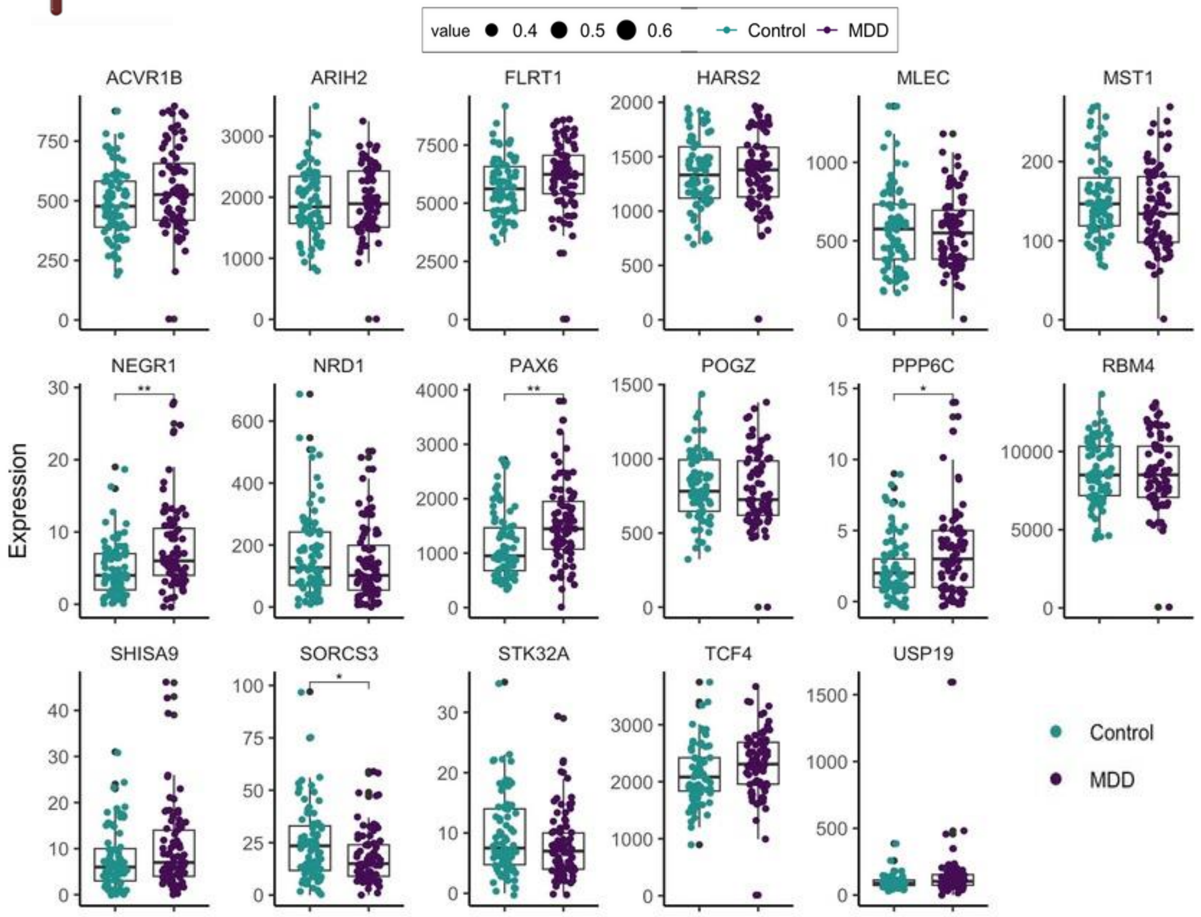


**Suppl Figure 3. FDR analysis of differentially expressed genes (DEGs) in PBMCs of MDD patients.** Box plots showing DEGs in the PBMCs (dataset: Trang T. L. et al., 2018) of MDD patients compared to healthy controls, after applying false discovery rate (FDR) correction for multiple comparisons.

### Suppl. figure 4

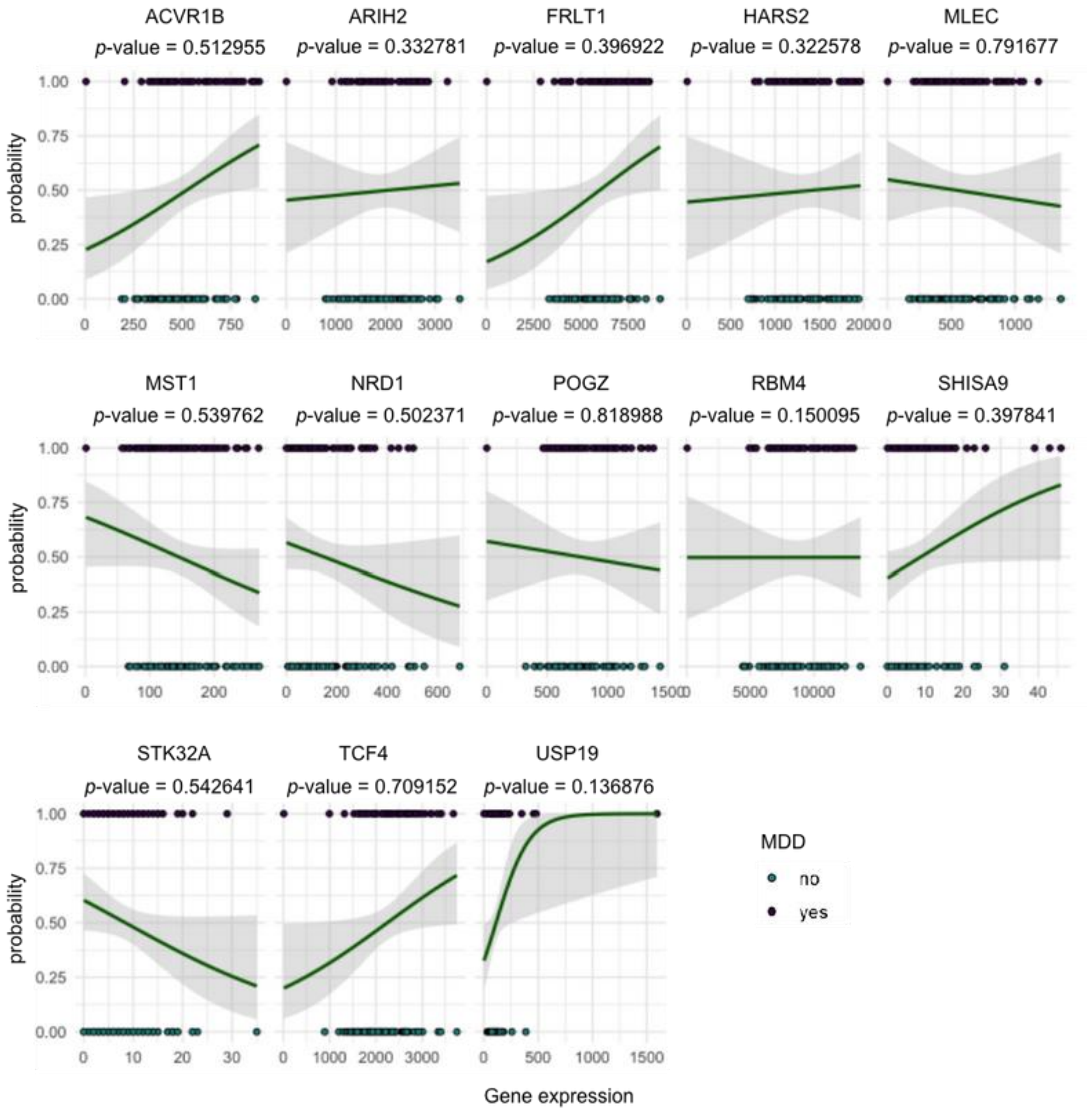


**Suppl. Figure 4. The probability of developing MDD based on the expression DEGs in PBMCs of MDD patients.** Scatter plots display the binomial logistic regression results for the expression of each DEG across the MDD group compared to healthy controls (dataset: Trang T. L. et al., 2018). The MDD groups are represented with green color for "yes" (1 = presence of the MDD) and gray color for "no" (0 = healthy controls).

a b

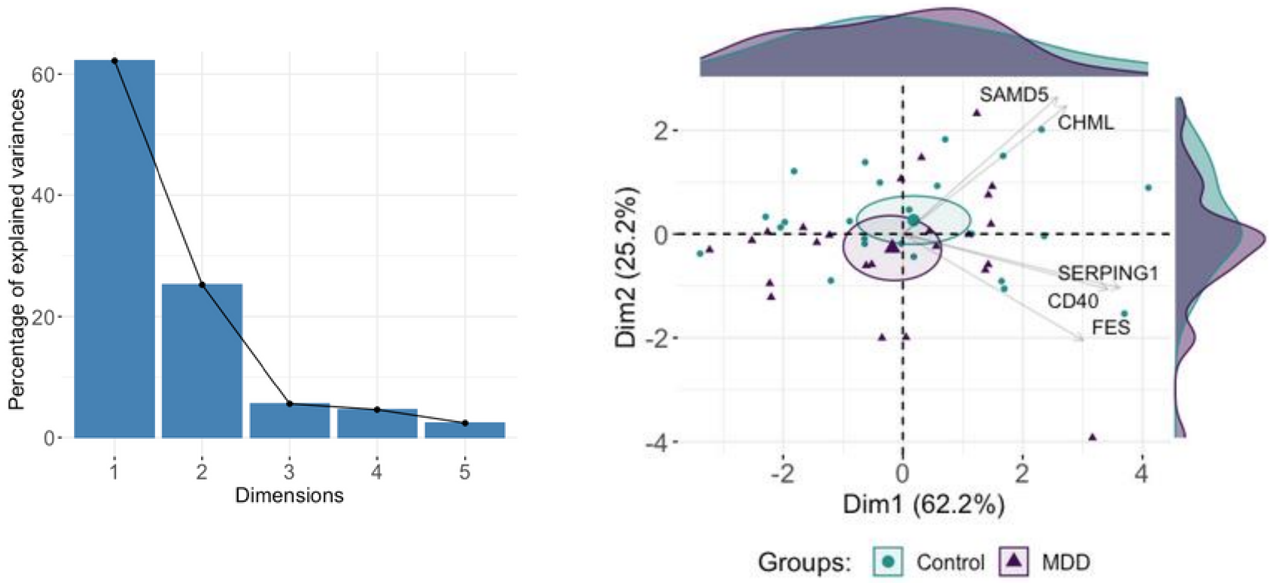


**Figure 4. PCA analysis of CAC analysis.** (**a**) Bar plot displaying the percentage of explained variance for each principal component (PC) dimension from a Principal Component Analysis (PCA) with spectral decomposition. (**b**) PCA plot illustrating the stratification of MDD patients and healthy controls based on PBMCs. Genes with positive correlations align on the same side of the plot, while negatively correlated genes point in the opposite direction. Small circles represent concentration ellipses around the mean points of each group. The accompanying histograms depict the density distribution of samples (individuals) for each group. (**c**) Heatmaps obtained from exploratory factor analysis of the 17 shared genes with negative and positive loadings visualized for MDD patients and health controls from the Tang T. L. et al. dataset. (**d**) Correlograms show the topological correlation pattern among shared genes. The color scale bar represents the range of Spearman's rank correlation coefficient for controls (left side of the graph) and MDD patients (right side).
